# Genetic modification of *Candida maltosa*, a nonpathogenic CTG species, reveals *EFG1* function

**DOI:** 10.1101/2023.11.08.566314

**Authors:** Marco Chávez-Tinoco, Luis F. García-Ortega, Eugenio Mancera

**Affiliations:** Departamento de Ingeniería Genética, Unidad Irapuato, Centro de Investigación y de Estudios Avanzados del Instituto Politécnico Nacional, Irapuato, Mexico

**Keywords:** CUG-Ser1 yeasts, CTG clade, *Candida maltosa*, pathogenic yeasts, nonpathogenic *Candida*, *EFG1*

## Abstract

*Candida maltosa* is closely related to important pathogenic *Candida* species, but it has been rarely isolated from humans. Therefore, through comparative studies, it could be a powerful model to understand the pathogenicity underpinnings of *Candida* species. To facilitate studying *C. maltosa* at a molecular level, we built a cohesive genomic sequence composed of 45 scaffolds, a substantial improvement from the thousands of contigs of the available draft. Comparison with *C. albicans* and *C. tropicalis* revealed a reduction in the total number of genes in *C. maltosa*. However, gene loss seems not to be associated to its avirulence given that most pathogenicity genes were also present in *C. maltosa*. To genetically edit *C. maltosa* we generated triple auxotrophic strains so that gene deletions can be performed as has been routinely done in pathogenic species. As a proof of concept, we generated gene knockouts of Efg1, a transcription regulator involved in filamentation and biofilm formation in pathogenic species. Although in *C. maltosa* Efg1 also played a role in these processes, it seems to rather be a repressor of filamentation. The genome assembly and auxotrophic mutants developed are a key step to start using *C. maltosa* for comparative studies at a molecular level.

## INTRODUCTION

Fungi from the CTG or CUG-Ser1 clade have attracted human attention for centuries given their medical relevance. This group of ascomycetous yeasts received their name because they translate the CUG codon to serine instead of leucine as most other eukaryotes do (Krassowski et al., 2018). Several of the clinically most important species of fungi such as *Candida albicans*, *C. tropicalis*, *C. parapsilosis* and *C. auris* belong to this clade (Butler et al., 2009; Jackson et al., 2009). Worldwide, these microorganisms are responsible for ∼700, 000 candidiasis cases every year, being an important part of the total number of infections caused by fungi and among the top health threats according to the World Health Organization (Bongomin et al., 2017; Parums, 2022). Interestingly, among the members of the CTG clade there are also species that have been very rarely isolated from humans. These species come from a variety of environments, including soil, air, water and other animals such as insects (Maciel et al., 2012; Mauersberger et al., 1996; Vadkertiová & Sláviková, 2006; Zhai et al., 2019). Several of the non-virulent species are closely related to pathogenic species (Figure 1) and given their phylogenetic position, the ability to colonize humans is thought to have originated repeated times within the clade (Gabaldón et al., 2016; Schikora-Tamarit & Gabaldón, 2022). Due to their phylogenetic proximity, non-virulent CTG species offer a valuable point of comparison to better understand the pathogenicity of these microorganisms.

**Figure 1.**
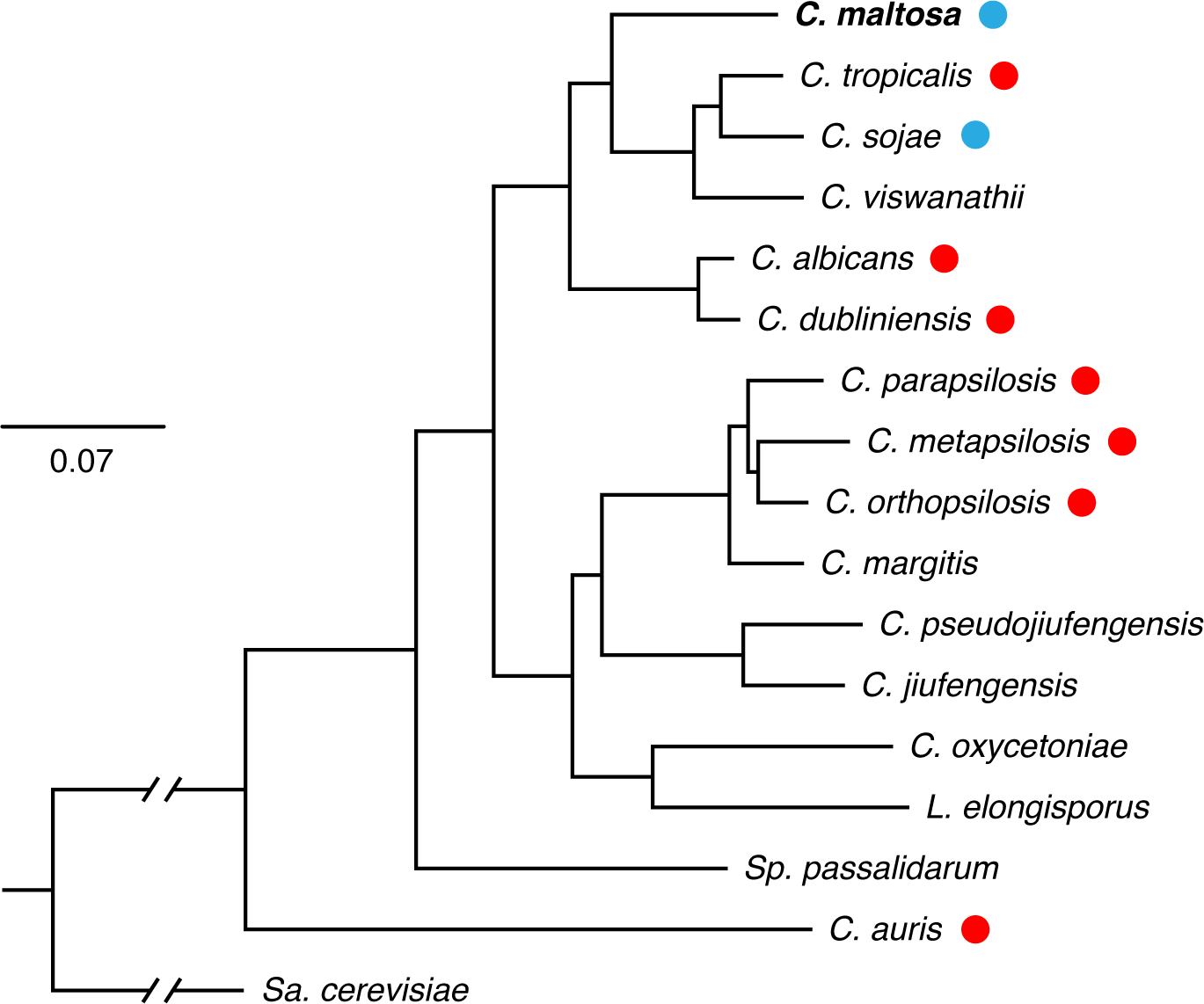
Phylogenetic relationship of *C. maltosa* with its closest CTG species and their association to humans. Maximum-likelihood phylogenetic tree based on the alignment of 569 single copy orthologs that were conserved in all the genomes of the species included (Materials and Methods). Numbers on the branches show bootstrap support based on 1,000 replicates and the genetic-distance scale represents substitutions per nucleotide. Red circles denote pathogenic species while blue circles non-pathogenic ones according to Gabaldon T. *et al*., (2016). *C. maltosa* was not included by Gabaldon T. *et al*., (2016) but given previous evidence and the frequency of isolation from humans it can be considered non-pathogenic.

Not surprisingly, most of the work to understand the molecular mechanisms that are responsible for pathogenicity has been targeted to the virulent species, mostly to *C. albicans.* Research in other virulent species has accelerated in recent years as some have become more common etiological agents. This is especially the case for *C. auris* which is an emerging pathogen of growing concern given its resistance to multiple of the typical antifungal drugs (Muñoz et al., 2018). On the other hand, considerably less is known about the biology of non-virulent or less virulent CTG species. Several studies have compared *C. dubliniensis* at a genomic and molecular level to *C. albicans,* since it is phylogenetically its closest relative and considerably less virulent (Flanagan et al., 2018; Jackson et al., 2009; Moran et al., 2012). However, *C. dubliniensis* is still considered an inhabitant of the human body. The genome sequences of several nonpathogenic CTG species are available (e.g., *Candida maltosa, C. sojae, C. tenuis, Debaryomyces hansenii* and *Meyerozyma caribbica*) and broad genomic comparisons have helped to generate hypothesis about the pathways involved in virulence (Butler et al., 2009). Research in non-pathogenic CTG species has also focused on biotechnological applications, but, overall, it is considerably little what we know about the ecology and adaptation to non-human environments of these species.

*C. maltosa* is a CTG species closely related to *C. tropicalis,* one of the most important human pathogens in the clade. It is also phylogenetically close to *C. albicans* and *C. parapsilosis*, which together with *C. tropicalis* account for most human infections caused by CTG species (Turner & Butler, 2014). The first strain of *C. maltosa* was described in 1964 coming from the adhesives of neutralizing tanks used to produce monosodium glutamate in Japan (Komagata & Katsuya, 1964). As far as we know, there are two reports of its isolation from immunocompromised patients and recently it was detected in human intrauterine samples through ITS amplicon sequencing (Liu et al., 2022; Mauersberger et al., 1996). However, the low frequency of isolation from humans suggests that the human body is not part of its core ecological niche, at least not more than it is for other non-pathogenic species such as the baker’s yeast. Furthermore, given the interest in employing *C. maltosa* for industrial processes, extensive testing in human cell cultures and laboratory animal models strongly indicated that it is a non-pathogenic species (Mauersberger et al., 1996). Strains of this species often come from industrial settings or from a variety of natural environments associated to anthropogenic activities, especially if enriched with hydrocarbons (Carmel Coronado et al., 2019; El-Latif Hesham et al., 2018). Historically it has received considerable attention due to its ability to grow in a variety of substrates such as carbohydrates, fatty acids and n-alkanes (Kawai et al., 1991; Mauersberger et al., 1996). For example, in the decade of 1980 it was intensively used in the former USSR to produce single-cell protein as animal fodder from n-alkanes in fuel-oil distillates (Gradova & Kovalskii, 1978; Mauersberger et al., 1996).

The industrial and biotechnological potential of *C. maltosa* motivated important research efforts to understand the biology of this yeast, especially during the 1990s and at the beginning of the 2000s. Hundreds of mutants were generated using physical and chemical mutagens, and several of its genes were cloned (Becher et al., 1994; Kawai et al., 1991; Nakazawa et al., 1997; Takaku et al., 2001). This culminated in host-vector systems that allowed its genetic manipulation despite the difficulties associated to the lack of a conventional sexual cycle that is typical of these fungi. However, only counted number of gene-mutants have been generated using these systems and the secondary effects of the physical and chemical mutagenesis in these strains remains unclear. More recent work has focused on the capacity of *C. maltosa* to metabolize xylose and the genes required for this pathway (Ake et al., 2023; Defosse et al., 2018; Lin et al., 2010). Since it was considered a potential strain for industrial xylitol and ethanol production, the genome of *C. maltosa* was sequenced in 2013 using Illumina short-read technology. This genomic draft is considerably fragmented, consisting of thousands of contigs, and there was no publication associated with the draft where the genomic characteristics and methodology used for the sequencing and annotation were described.

Despite the attention that *C. maltosa* attracted at the end of last century given its biotechnological potential, it has been mostly neglected as a comparative model to understand the pathogenicity of its closely related CTG species. Here, we re-sequenced the genome of *C. maltosa* using long-read technology, assembling a much more cohesive genomic draft and complete gene annotation. We also generated a set of strains and cassettes that allow its rapid genetic modification. As proof of concept, we deleted both alleles of the *EFG1* gene which encodes a transcription regulator that controls processes associated to the virulence of closely related species. Even though *C. maltosa* is a nonpathogenic species inhabiting considerably different environments, characterization of the mutants showed that Efg1 is also involved in filamentation and biofilm formation. However, this transcription regulator seems to be a repressor of filamentation in this species. The genomic draft and genetic engineering tools developed here will allow using *C. maltosa* as a nonpathogenic reference to understand the molecular mechanisms implicated in the virulence of this medically important group of fungi. In addition, these tools will facilitate the use of this species for biotechnological applications.

## MATERIALS AND METHODS

### Genome sequencing and assembly

High molecular weight genomic DNA was extracted from an overnight YPD culture of *C. maltosa* Xu316 (Lin et al., 2010) using a QIAGEN Genomic-tip 20/G according to the manufacturer’s instructions. Genomic DNA was sequenced in parallel using the PacBio Sequel II platform and the DNBSeq platform at Novogene (Beijing, China) and BGI (Shenzhen, China), respectively. For PacBio long reads, three single-molecule real-time cells were used to generate a total of 586,673 raw reads with a length of between 30 and 52 kb and an estimated N50 value of 76.5 kb. Low-quality reads were filtered using Filtlong (v0.2.0) with the parameters “keep percent 80” and “length weight 10” (Wick & Menzel, 2009). For the DNBSeq PE150 short reads, over 8 million raw reads were filtered by fastp (v0.23.2) using −c −x − y options for polyX trimming in 3′ ends, base correction in overlapped regions, and low complexity filtering, respectively (Chen et al., 2018). The preliminary assembly was obtained using CANU (v1.8) with the high quality PacBio reads (Koren et al., 2017). The clean DNBseq reads were then used in five consecutive rounds to polish the assembly with Pilon (v1.23) (Walker et al., 2014). Genome completeness was evaluated with BUSCO v5.4.3(Simão et al., 2015) using the lineage data set saccharomycetes_odb10.

### Gene annotation

Gene annotation of the *C. maltosa* and *C. soja* genomes was performed using the MAKER pipeline (v2.31.8) (Campbell et al., 2014). This included an initial round of evidence-based alignment followed by two rounds of training and prediction using the SNAP and Augustus gene prediction programs (Korf, 2004; Stanke et al., 2006). In the first round, we built a pangenome from the genomes of *C. albicans*, *C. tropicalis* and *C. parapsilosis*, and the resulting set of transcripts and proteins was provided as EST and protein evidence, respectively. In this initial round, as masking evidence, we also provided a custom repeat library generated with RepeatMasker 4.1.5 and RepeatModeler 2.0.4 (Smit A et al., 2013). The resulting gene models with proteins longer than 50 residues and AED scores below 0.25 were retained as evidence to train SNAP and Augustus for the two additional rounds of gene prediction. Augustus was trained with BUSCO using the initial HMM model of the fungi odb10 dataset. The functional annotation was performed using a BLAST homology search for each predicted protein sequence against the Uniprot database. Only the hits with an e-value inferior to 10E-6 were considered. Finally, InterProscan (v5.35.74) was used to detect protein domains, Gene Ontology terms and KEGG mappings from the InterPro database (Jones et al., 2014).

### Genomic comparison to closely related species

To compare the genome of *C. maltosa* with the ones of its closest related species, *C tropicalis, C. sojae*, and *C albicans,* we followed the workflow described by (Paril et al., 2023). In short, this consisted in running Orthofinder (v2.5.5) (Emms & Kelly, 2019) using the proteomes of the four species and using HMMER3 and Panther HMMs protein-coding gene family models (Mistry et al., 2013; Thomas et al., 2021). To estimate significant gene family expansions or contractions across genomes we used CAFÉ (v5) (Bie et al., 2006). For phylogenetic reconstruction, we ran Orthofinder with the proteomes generated here of *C. maltosa* and *C. sojae*, together with the 11 closest *Candida* species with available genomic annotations at GenBank, representatives of other CTG subclades (*Spathaspora passalidarum* and *Lodderomyces elongisporus)*, *C. auris* as an important pathogen and *S. cerevisiae* as an outgroup. The 569 single copy orthologs that were conserved in all the seventeen genomes were aligned using MUSCLE (v3.8.31) (Edgar, 2004) and the alignments were polished using TrimAl (v1.2) (Capella-Gutiérrez et al., 2009). Finally, the concatenated alignments were used to reconstruct the phylogeny using RAxML-HPC2 on XSEDE (v8.2.12) in CIPRES (Miller et al., 2015) with the PROTCATWAG model and a total of 1000 bootstrap replicates(Muñoz et al., 2018).

### Identification of genes associated with pathogenicity

Genes associated with pathogenicity were obtained from previously curated catalogs in *C. albicans* (33 genes from Gómez-Gaviria et al., 2023, and 674 from Noble et al., 2010) resulting in a set 629 genes. ALS, TLO and SAP genes were obtained from Jackson et al., (2009) and Oh et al., (2019). Genes related to filamentous growth were identified from the Candida Genome Database (http://www.candidagenome.org/) using the phenotypes section in tools utility, choosing the Phenotype Term “filamentous growth” and including the following phenotypes: “filamentous growth: abnormal”, “filamentous growth: increased”, “filamentous growth: decreased”, “filamentous growth: absent”, “filamentous growth: decreased rate” and “filamentous growth: delayed”. Orthologs between *C. albicans*, *C. tropicalis*, *C. maltosa* and *C. sojae* of the sets of genes mentioned above were defined from the genome-wide orthology assignments established before with Orthofinder. To verify the absence of pathogenicity-associated genes not found in *C. maltosa,* we conducted pseudogene predictions using Pseudopipe (Z. Zhang et al., 2006) on the genomic draft with default parameters, and the protein sequences of *C. albicans* as queries. Putative pseudogenes were filtered by excluding any matches that overlapped with functional gene annotations, transposon elements, or sequences shorter than 150 bp.

### Generation of auxotrophic strains

Given that *C. maltosa* is closely related to *C. tropicalis,* auxotrophic strains were generated using the SAT1-flipping strategy that has been optimized for *C. tropicalis* (Mancera et al., 2019). In brief, deletion cassettes were generated by PCR with 99 pb primers that have ∼70 base sequences that are identical to the 5’ and 3’ of the genes *HIS1*, *LEU2* and *ARG4* of *C. maltosa* (Supplementary Table 2). Plasmid pEM018 was used as a template. The resulting cassette was purified with a QIAGEN MinElute PCR Purification Kit and ∼0.5 µg was transformed by electroporation as previously described (Mancera et al., 2019). The *C. maltosa* strain transformed was Xu316 (Lin et al., 2010) (Table 1), strain for which a genome draft was already available. Colony selection was done in YPD medium plates (2% yeast extract, 2% peptone, 2% glucose and 4% agar) containing 400 µg/ml of nourseothricin (NAT) and the correct integration of the SAT1 cassette was verified by colony PCR of the 5’ and 3’ flanks. The deletion of the second allele was also verified by the absence of the corresponding ORF through colony PCR and by the inability of the strains to grow in SD media lacking the corresponding amino acid. To recycle the SAT1 marker, previously to the second round of transformation, the cassette was flipped out by growing cell overnight at 30 °C in liquid YNB medium with 2% casamino acids (Mancera et al., 2019). Flipped cells were subsequently screened for the loss of NAT resistance by replica plating in YPD medium plates supplemented with 400 µg/ml NAT.

**Table 1.** Comparison of the genomic characteristics of the current (2023) and previous (2013) *C. maltosa* genome assemblies.

| Genomic features | 2023 Assembly | 2013 Assembly |
| --- | --- | --- |
| Number of scaffolds | 45 | 2,947 |
| Total size of scaffolds (bp) | 14,040,896 | 12,826,280 |
| Total scaffold length as % of assumed genome size | 100 | 88.45 |
| N50 (bp) | 1,315,209 | 11,156 |
| Number of genes | 5,518 | 5,986 |
| Average gene length (bp) | 1,597 | 1,715 |
| Number of exons | 8,534 | 7,042 |
| BUSCO assembly completeness | 95.2% | 94.60% |
| BUSCO annotation completeness | 94.80% | 89.80% |

### Reintegration of nutritional markers and deletion of target genes

To generate reference strains, we reintegrated the *C. albicans HIS1*, *LEU2* and *ARG4* markers into the triple auxotrophic strains, similarly to what has been done for the deletion of the two alleles of a gene of interest (Mancera et al., 2019; Noble & Johnson, 2005). The reintegration cassettes were generated by fusion PCR of three fragments, ∼300 bp sequences flanking the *C. maltosa* genes and the sequence coding for the *C. albicans* nutritional marker. The primers used are listed in Supplementary Table 2. As a template to amplify the flanks we used genomic DNA of *C. maltosa,* and for the markers we employed plasmids pEM001, pEM002 and pEM00 3 for *HIS1, LEU2*, and *ARG4*, respectively (Supplementary Table 2) (Mancera et al., 2019). The resulting cassettes were purified and concentrated using a QIAGEN MinElute PCR Purification Kit and ∼0.5 µg was transformed by electroporation. Selection of transformants was performed by growth on SD plates without the corresponding amino acid. The integration in the correct locus was then verified by colony PCR of the 5’ and 3’ flanks. This resulted in the reintegration of the *C. albicans HIS1, LEU2* and *ARG4* nutritional markers in *C. maltosa* at the corresponding loci. The presence of the markers was finally verified by colony PCR with primers that target a region within the marker genes. The same strategy was followed for the deletion of the two alleles of *EFG1*. *C. albicans HIS1* nutritional marker was used to delete the first allele, and *LEU2* to delete the second, as has also been previously described (Mancera et al., 2019; Noble & Johnson, 2005).

### Filamentation essays

Filamentation was assessed as previously described (Lackey et al., 2013). Briefly, an overnight culture in YPD at 30 °C was washed twice with water and was subsequently resuspended in the inducing medium to a final OD_600_ of 1.5. The following media were tested: minimum medium (0.17g Yeast Nitrogen Base without amino acids and ammonium sulfate, 0.5g ammonium sulfate dibasic, 2g glucose) adjusted at three different pH, 4.5, 6.0 and 7.0, RPMI medium (MP biomedicals, added with sodium bicarbonate and 2% glucose), Lee medium (Lee et al., 1975; Q. Zhang et al., 2015), Synthetic dextrose medium (SD) (Naseem et al., 2020), and synthetic defined medium supplemented with 0.75% glucose and 50% fetal bovine serum (FBS) (Lackey et al., 2013; Mancera et al., 2015). Cells were inspected under an optical microscope with phase contrast at the time of induction and after 0, 3 and 6 hours of incubation at 37 °C shaking in a roller drum. Micrographs were obtained using a Leica DMRX microscope using DIC and fluorescence of cells stained with calcofluor as previously described (Ramos et al., 1999).

### *In vitro* determination of biofilm formation

To assess biofilm formation, we followed the protocol previously described for other CTG species (Mancera et al., 2021; Nobile et al., 2012). In brief, non-tissue culture treated six-well polystyrene plates were preincubated with adult bovine serum (BSA) overnight at 37 °C and shaking at 200 rpm. After removing the BSA and washing with phosphate buffered saline (PBS), 4 mL of fresh media were inoculated to an OD_600_ of 0.5 with cells from an overnight YPD culture. For adhesion, cells were incubated shaking at 200rpm and 37 °C for 90 minutes. Non-adhered cells were then aspirated, and the wells were washed twice with PBS to finally add 4 mL of the medium where biofilm formation would be assessed. After 48 hrs., unadhered cells were removed and biofilms were visually inspected. To determine the medium where *C. maltosa* formed thicker biofilms, we tested the following media: Lee medium (Lee et al., 1975; Q. Zhang et al., 2015), Lee medium supplemented with 1.25% glucose, Spider medium with 1% mannitol, glucose or xylose as carbon source, cornmeal liquid medium (prepared from cornmeal medium in cold water and filtering insoluble components before autoclaving) (Gupta et al., 2019), and minimum media at pH 4.5. Based on the qualitative inspection of biofilms in the different media, we employed Spider with 1% glucose to determine biomass dry weight of the *efg1* mutant. This was done by scraping and aspirating the 48 hrs. biofilms onto a cellulose filter paper. Weight was determined in an analytical balance after drying biofilms for 24 hrs. and subtracting the weight of a control well where cells had not been inoculated. Five replicate biofilms grown in separate wells were used for each strain.

## RESULTS

### A cohesive *C. maltosa* genome assembly for comparative genomics and genetic engineering

As mentioned before, the currently available genomic sequence of *C. maltosa* was sequenced in 2013 using short-read technology and the eight pairs of expected chromosomes are divided in over 2,500 contigs. The considerably fragmented status of this genome draft makes it difficult to perform thorough genome comparisons and design efficient genetic modification experiments. To achieve a more cohesive genome draft of *C. maltosa* we performed long and short read sequencing using PacBio and DNBSeq technologies, respectively. Using the long reads, we tested three different genome assemblers (Canu, Falcon and Smartdenovo) to generate a new genome assembly. Based on the N50, L50 and NG50, the assembly made by Canu was selected and subsequently polished using the PE150 short reads. The resulting genome draft is composed of 45 contigs with a N50 of 1,315,209 bp. The size of the new genome assembly (14.04 Mb) was considerably larger than that of the previous *C. maltosa* draft (12.8 Mb), while the GC content is very similar (∼34%). In addition, genome size and GC content were comparable to those of closely related species (*C tropicalis* 14.3 Mb, 33% GC, *C. sojae*, 15.1 Mb, 32% GC and *C albicans*, 14.3 Mb, 33% GC). To assess completeness, we performed BUSCO analysis showing a coverage of 95.2% that represents an improvement of almost 1% from the previously available assembly (Table 1).

To annotate the genes in the new *C. maltosa* genome assembly we built a reference pangenome from the genomic annotations of the three closest CTG species and whose genomes are well annotated*, C tropicalis, C albicans* and *C parapsilosis*. Using this reference set of genes increased the possibility of identifying genes in the new genome by reducing the genes that are not present in the reference set. This approach predicted a total of 5,518 genes composed of 8,534 exons in the *C. maltosa* assembly (Table 1). The average gene length is 1,597 bp and the average number of exons per gene is 1.5. BUSCO analysis of the annotated transcriptome using the Saccharomycota database, showed a coverage of 94.8%, which represented an evident improvement from the annotation of the previously reported assembly (89.8%) (Table 1). In contrast, when we only used the *C, tropicalis* genome as a reference gene set for the annotation, we found 5,316 gene models and the BUSCO completeness went down to 89.6 %.

The genome of *C. sojae* has been recently sequenced using long read technology and it is almost assembled to the chromosome level (Guin et al., 2020). However, there is no publicly available gene annotation that could be used to compare the gene content to other species, such as *C. tropicalis* or *C. maltosa*. Taking advantage of the pangenome built for the annotation of *C. maltosa* we also predicted gene models for the *C. sojae* genome. The resulting annotation is composed of 6,216 genes which is very close to the 6,441 genes of the latest *C. tropicalis* annotation. The genome of *C. maltosa* on the other side, had close to seven and nine hundred genes less than these genomes, respectively.

### Genes associated to pathogenicity are present in *C. maltosa*

To identify genes that could be responsible for the differences in virulence between *C. maltosa* and closely related pathogenic *Candida* species, we first defined orthologous genes throughout the genome using OrthoFinder. We considered *C. tropicalis* as the closest species to *C. maltosa* that is human associated, and *C. albicans* as the best studied and most virulent species of the CTG clade (Figure 1). *C. sojae* was also included in the analysis since it is as closely related to *C. maltosa* as *C. tropicalis*, but it is considered not pathogenic (Figure 1). As mentioned above, we needed to build the gene models for *C. sojae* since there was no publicly available annotation. We identified 4,594 orthogroups that are shared between all four species from a total of 5,731, and most of them are single copy orthogroups. *C. tropicalis* had the most unassigned genes (303), followed by *C. albicans* (194), *C. sojae* (111) and *C. maltosa* (61) (Figure 2). In agreement with its smaller genome, *C. maltose* had the largest number of gene family contractions between these species (34 contractions).

**Figure 2.**
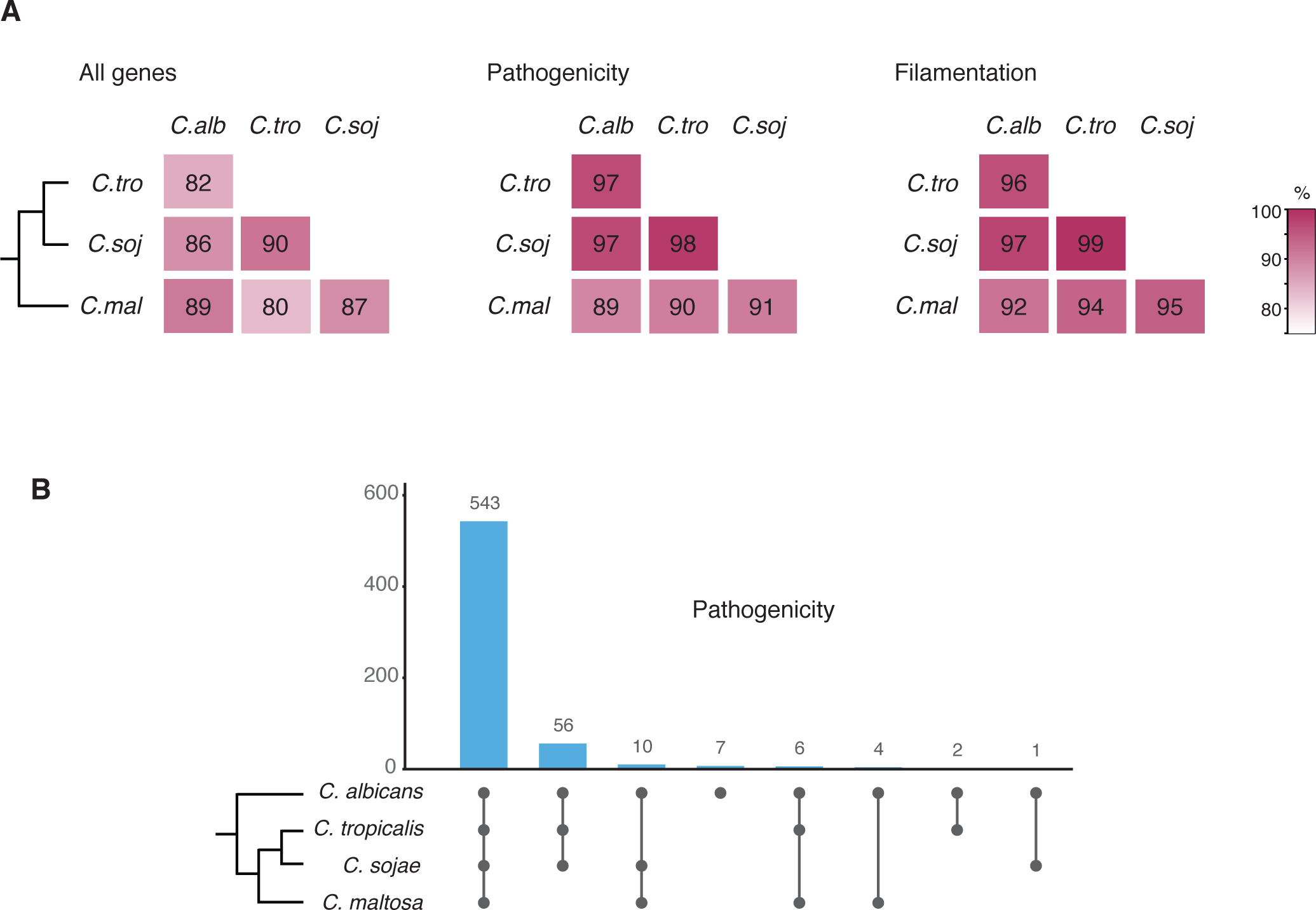
Most genes associated to pathogenicity are present in *C. maltosa.* A) pair-wise comparison of shared orthologue genes between *C. maltosa* and its closest relatives. Each of the three heatmaps shows, from left to right, overall genes in the genome, pathogenicity associated genes and genes related to filamentation (Material and Methods). B) bar plot showing the shared pathogenicity genes between the same four species, but in all possible comparisons. Most genes (86%) are conserved in the four species analyzed.

To identify pathogenic associated genes among the set of orthologs in these species we used two previously curated gene catalogs (Gómez-Gaviria et al., 2023; Noble et al., 2010). These lists include 629 genes of *C. albicans* that have been implicated in different aspects of pathogenicity such as hyphae and biofilm formation, adhesion, evasion of the immune system and invasion of the host (Supplementary Table 2). Of these genes, 96.6% have an ortholog in *C. tropicalis*, 89.8% in *C. maltosa* and 96.9% in *C. sojae* (Figure 2). The percentage of shared genes in this category is more or equal to the proportion of overall paralogs between *C. albicans* and these three species (82%, 89%, and 86%, respectively) and is more than what would be expected by chance (< 30%).

We also assessed gene conservation in other specific gene families that have been linked to virulence in these species, namely telomere-associated genes (*TLOs*), agglutinin like-sequences (*ALSs*), secreted aspartyl proteinases (*SAPs*) and filamentation genes (Table 2). TLOs are a family of genes that have a conserved N-terminal Med2 domain and therefore, apart from their role in virulence, are thought to be related to general transcription (Flanagan et al., 2018; Haran et al., 2014). *C albicans* has 15 *TL*O genes (Jackson et al., 2009), while only one was present in *C. maltosa.* However, *C. tropicalis* and *C. sojae* also had only one *TLO* ortholog, suggesting that the ancestor of these three species had most probably also one and the number is not related to their ability or lack thereof to colonize humans. The *ALS* genes are central for cell adhesion, tissue invasion and biofilm formation (S.-H. Oh et al., 2019; S. H. Oh et al., 2021). *C. albicans* is known to have nine of these genes (Jackson et al., 2009) and 16 have been found in *C. tropicalis* (S. H. Oh et al., 2021). We found eleven *ALS* orthologs in *C. maltosa* and 13 in *C. sojae*. *SAPs* have also been associated to adhesion but their role in virulence is most probably related to host tissue invasion through their hydrolytic activity (Parra-Ortega et al., 2009; Taylor et al., 2005). *C. albicans* has ten different *SAPs* and *C. tropicalis* six (Jackson et al., 2009), while the species not associated to humans had ten and six genes in the case of *C. maltosa* and *C. sojae,* respectively (Table 2). In addition, given the importance of filamentous growth for the biology of these species, a set of 447 genes related to hypha formation in *C. albicans* according to CGD was used to identify orthologous genes in *C. maltose* and related species. Almost all these genes have an ortholog in the other three species, although the set is overall less conserved in *C. maltosa* (96.2% in *C. tropicalis*, 97.3% in *C. sojae* and 92.0% in *C. maltosa*) (Figure 2).

**Table 2.** Number of *ALS*, *TLO* and *SAP* genes in *C. maltosa* and its three closely related species.

| Species | ALS | SAP | TLO |
| --- | --- | --- | --- |
| *C. albicans* | 9 | 10 | 15 |
| *C. maltosa* | 11 | 10 | 1 |
| *C. sojae* | 13 | 6 | 1 |
| *C. tropicalis* | 16 | 6 | 1 |

To verify that the pathogenicity and filamentation associated genes (seven and nine genes, respectively) detected as missing in *C. maltosa* are absent from the genome, we searched for vestiges of these ORFs employing a pipeline to detect pseudogenes (See Material and Methods). Of the16 genes, only two were identified as a pseudogene, strongly suggesting that the observed absences are actual gene losses.

### A set of *C. maltosa* auxotrophic strains for its genetic modification

As most of its closely related species, *C. maltosa* has only been isolated in a diploid state and it has not been observed to undergo a sexual cycle. For these reasons, to generate a gene knockout strain it is necessary to directly delete the two alleles of the gene of interest. To this end we generated auxotrophic strains for three nutritional markers in *C. maltosa*, leucine, histidine, and arginine. Two of the markers can be used to delete the two alleles of the targeted gene and the third for reintegration of a copy for complementation assays. This strategy has been routinely used in other related *Candida* species, even for the generation of large collections of knockout strains (Holland et al., 2014; Noble et al., 2010). In addition, auxotrophic strains have been instrumental in implementing CRISPR based strategies in related species (Nguyen et al., 2017). To generate the triple auxotrophic strains, we tandemly deleted the two alleles of the genes *LEU2*, *HIS1* and *ARG4* using the SAT-1 flipping system that has been optimized for *C. tropicalis* (Mancera et al., 2019). The flipping strategy was an effective method to genetically modify *C. maltosa*, although considerably laborious given that the *SAT1* marker must be flipped out between every round of allele deletion and due to the incubation time needed for the cells to express the resistance marker before selection can be applied. These factors can be a considerable disadvantage for constructing collections of mutant strains when compared to the strategy that uses multiple nutritional markers (Mancera et al., 2019; Noble & Johnson, 2005).

The ORFs that corresponded to the *LEU2*, *HIS1* and *ARG4* genes in *C. maltosa* were identified as the best BLAST reciprocal hits to the *C. tropicalis* and *C. albicans* genes. The protein sequence identity of the three genes among the three species ranged from 91.69 to 95.44% and at the nucleotide level it went from 84 to 85%. The differences at the nucleotide level permitted using the genes of *C. albicans* as the nutritional markers so that they are not inserted at the endogenous *LEU2*, *HIS1* and *ARG4* loci, but rather at the locus that is being targeted for deletion. On the other side, the conservation at the protein level ensured functional complementation by the *C. albicans* markers. Two independent triple auxotrophic strains were generated in the genetic background for which a genomic sequence was previously available, Xu316 (Lin et al., 2010) (Table 3). In addition, in the process of generating these strains, single leucine auxotrophic strains and double leucine and histidine auxotrophic strains were generated (Table 3). All these strains are a valuable resource for the genetic modification of *C. maltosa*.

**Table 3.** *Candida* strains used and generated in this study.

| Name | Species | Background | Genotype | Source |
| --- | --- | --- | --- | --- |
| Xu316 | *C. maltosa* | Xu316 | *WT* | Lin *et al.*, 2010 |
| CMMC14 | *C. maltosa* | Xu316 | *leu2Δ::FRT/leu2Δ::FRT* | This Study |
| CMMC15 | *C. maltosa* | Xu 316 | *leu2Δ::FRT/leu2Δ::FRT* | This Study |
| CMMC27 | *C. maltosa* | Xu316 | *his1Δ::FRT/his1Δ::FRT leu2Δ::FRT/leu2Δ::* | This Study |
| CMMC28 | *C. maltosa* | Xu316 | *his1Δ::FRT/his1Δ::FRT leu2Δ::FRT/leu2Δ::* | This Study |
| CMMC45 | *C. maltosa* | Xu316 | *his1Δ::FRT/his1Δ::FRT leu2Δ::FRT/leu2Δ::FRT*<br>*arg4::FRT/arg4::FRT* | This Study |
| CMMC46 | *C. maltosa* | Xu316 | *his1Δ::FRT/his1Δ::FRT leu2Δ::FRT/leu2Δ::FRT*<br>*arg4::FRT/arg4::FRT* | This Study |
| CMMC66 | *C. maltosa* | Xu316 | *his1Δ::FRT/his1Δ::FRT LEU2 Calb/leu2Δ::FRT*<br>*arg4::FRT/arg4::FRT* | This Study |
| CMMC67 | *C. maltosa* | Xu316 | *HIS1 Calb/his1Δ::FRT LEU2 Calb /leu2Δ::FRT*<br>*arg4::FRT/arg4::FRT* | This Study |
| CMMC55 | *C. maltosa* | Xu316 | *his1Δ::FRT/his1Δ::FRT leu2Δ::FRT/leu2Δ::FRT*<br>*arg4::FRT/arg4::FRT efg1Δ:: Calb HIS1/efg1Δ:: Calb*<br>*LEU2* | This Study |
| CMMC56 | *C. maltosa* | Xu316 | *his1Δ::FRT/his1Δ::FRT leu2Δ::FRT/leu2Δ::FRT*<br>*arg4::FRT/arg4::FRT efg1Δ:: Calb HIS1/efg1Δ:: Calb*<br>*LEU2* | This Study |

### Efficient generation of knockout mutants through reintegration of nutritional markers

To test the gene knockout strategy, we deleted the ortholog of the gene encoding for the transcription regulator Efg1. In closely related species, Efg1 regulates several morphological transitions (REFs), but it is not an essential gene, being an ideal candidate for the proof of concept. The ORF of *EFG1* in *C. maltosa* was identified during the annotation of the genomic sequence and it was independently verified as the best reciprocal BLAST hit with the gene of *C. tropicalis*. For its deletion, *C. albicans LEU*2 and *HIS1* cassettes with ∼300 bp flanks of sequence identical to the up- and down-stream regions of the *C. maltosa EFG1* ORF were constructed by fusion PCR. These cassettes were used to tandemly delete the two alleles of *EFG1,* and the knockout was successfully verified by colony PCR with primers that annealed at the junctions of the cassettes and at the ORF itself. Our results showed that the *LEU*2 and *HIS1* genes of *C. albicans* can functionally complement the corresponding auxotrophies in *C. maltosa,* and that the strategy is an efficient way to generate gene knockouts as has been done in other *Candida* species (Mancera et al., 2019; Noble & Johnson, 2005).

To generate a set of reference strains we also reintegrated the amino acid markers of *C. albicans* at the endogenous *LEU2*, *HIS1* and *ARG4* loci (Table 2). To this end, similar deletion cassettes were constructed as for the deletion of *EFG1*, but the flanks of the cassettes were sequences identical to the up- and down-stream region of the amino acid genes. Growth of colonies in medium lacking the corresponding amino acid was used to select for transformants showing that *ARG4* from *C. albicans* is also able to functional complement the auxotrphy in *C. maltosa*. The correct integration of the markers was finally verified by colony PCR with primers directed to the junctions of the markers and the regions surrounding the *LEU2*, *HIS1* and *ARG4* loci. The resulting set of strains (Table 2) are the ideal references for the phenotypic characterization of knockout mutants generated using the genetic modification strategy implemented.

### *EFG1* has a role in filamentation and biofilm formation in *C. maltosa*

To assess whether the function of the transcription regulator Efg1 is conserved in *C. maltosa,* we tested the gene deletion mutant for its ability to filament and form biofilms. These two morphological transitions depend on this transcription regulator in *C. tropicalis*, *C. albicans* and other related species (Mancera et al., 2015, 2021). To this end, standard *in vitro* filamentation and biofilm formation assays were employed, although a variety of media were tested to identify the optimal condition for this species. For filamentation, minimum medium had been previously used for *C. maltosa* (Takaku et al., 2002) and it was where we observed that it filamented the most. However, despite the six different media tested, *C. maltosa* did not filament as efficiently as *C. tropicalis* and *C. albicans* in liquid cultures. When assessing the *efg1* mutant, surprisingly, we observed a 30% increase in filament formation after six hours in the inducing conditions compared with de reference strain (Figure 3). This filamentation increase contrasted the phenotype observed in *C. tropicalis*, in which the *efg1* mutant did not filament under the conditions tested. However, the deletion of *EFG1* has been previously associated to increased pseudohyphal growth in *C. parapsilosis* (Connolly et al., 2013).

**Figure 3.**
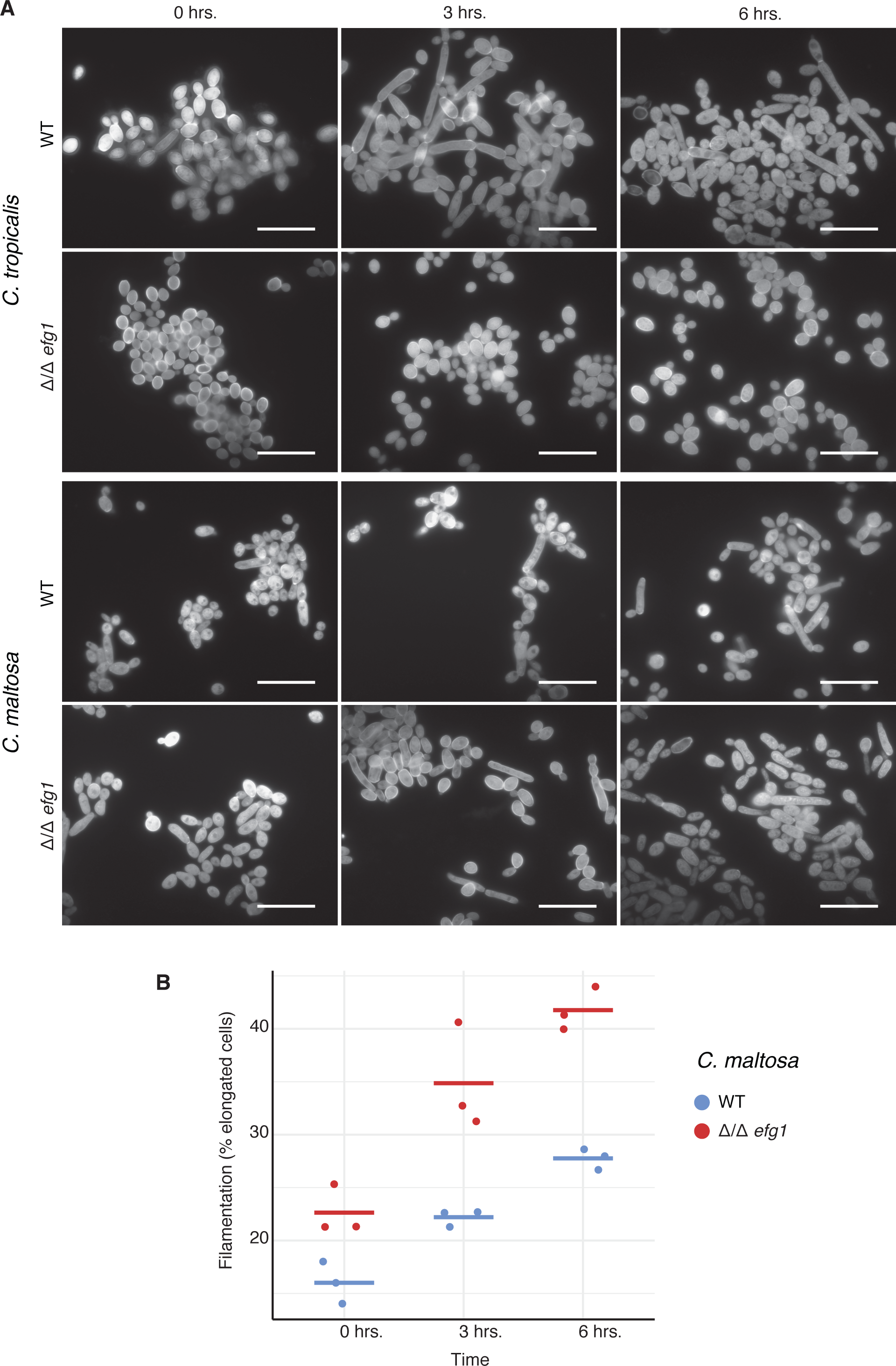
*EFG1* is involved in filamentation in *C. maltosa.* A) micrographs of *C. maltosa* and *C. tropicalis* with its Δ*/*Δ*efg1* mutants under filamentation inducing conditions (minimum media pH 4.5). Representative micrographs are shown at the time of transfer to the induction medium (0 hrs.) and after 3 and 6h of culture in these conditions. Scale bar represents 20 μm. B) quantification of the fraction of filamented cells of *C. maltosa* after transfer to inducing conditions through time.

**Figure 4.**
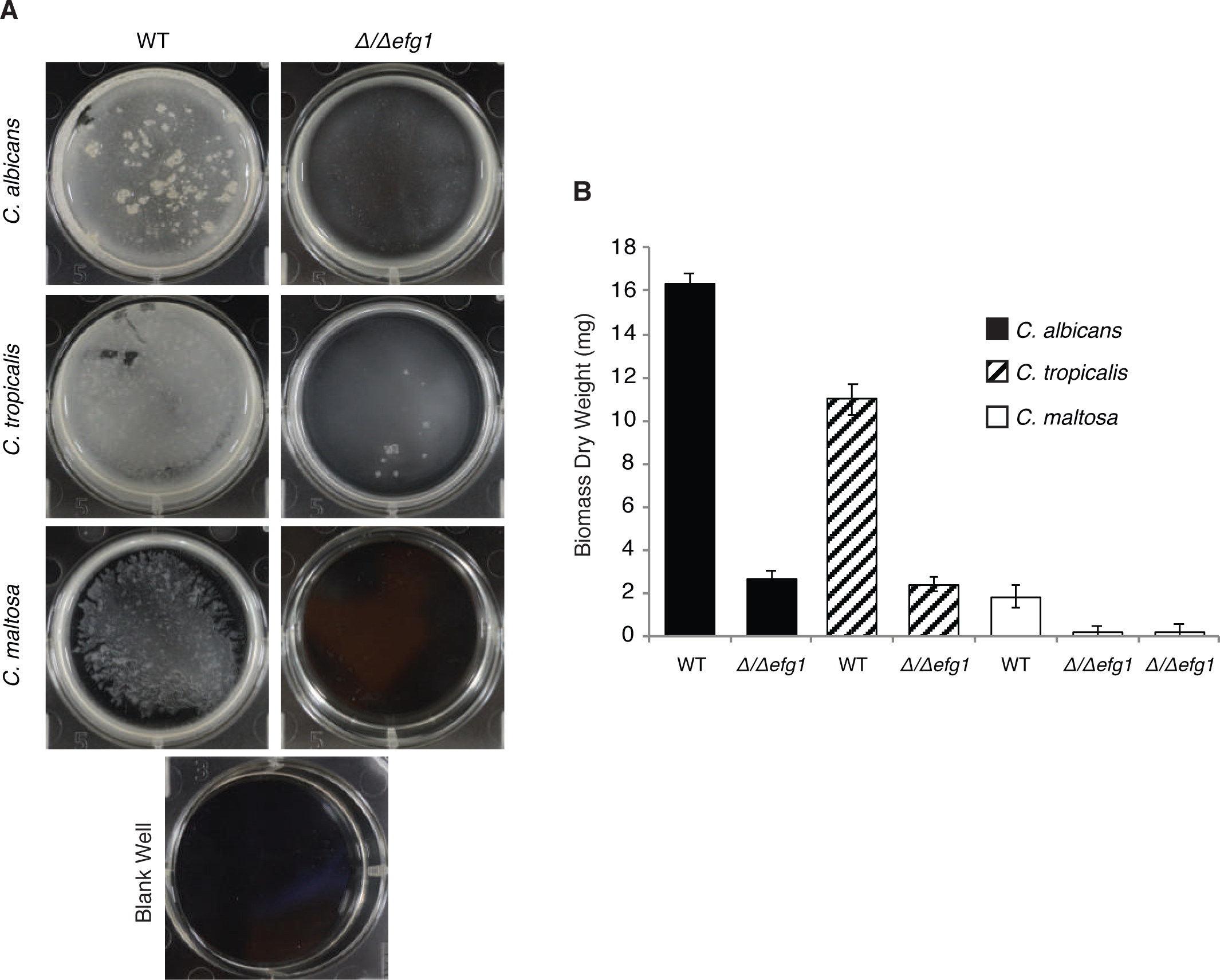
*EFG1* is required for biofilm formation in *C. maltosa.* A) biofilm formation by *C. maltosa, C. albicans* and *C. tropicalis,* and their corresponding Δ*/*Δ*efg1* mutants at the bottom of six-well polystyrene plates in Spider glucose medium (Material and methods). One well is shown per strain. B) comparison of the biomass dry weight of the biofilms. Dry wight of five replicates was used to determine the standard deviation.

For biofilm formation, performing qualitative assays we determined that Spider medium or Spider replacing mannitol by glucose as a carbon source, were the media where *C. maltosa* was better able to form biofilms. However, as for filamentation, in all media tested, the biofilms formed by *C. maltosa* were considerably thinner than those formed by *C. tropicalis* or *C. albicans*. To test biofilm formation in the *efg1* mutant we employed Spider with glucose since this medium has been shown to also be optimal for the sister species *C. tropicalis* and would therefore facilitate comparisons (Mancera et al., 2021). Although the biofilms formed by wildtype *C. maltosa* are relatively thin, we did observe a close to 90% reduction in the dry-weight biomass of the biofilms formed by the *efg1* strain (Figure 3). We also assessed biofilm formation in minimum medium since it was where this species filamented more efficiently. The results showed a similar trend, the *efg1* mutant formed considerably thinner biofilms and overall *C. maltosa* formed thinner biofilms than *C. tropicalis* and *C. albicans* (Figure 3). Together with the filamentation phenotypes observed, these results showed that the overall function of Efg1 as a morphological regulator is conserved in *C. maltosa*.

## DISCUSSION

Due to its phylogenetic position and avirulence, *C. maltosa* is a promising species to understand the mechanisms that underly the pathogenicity of closely related *Candida* species through comparative studies. Here, we generated a comprehensive genomic sequence and constructed a set of strains that will facilitate the genetic manipulation of *C. maltosa*. Analysis of the genomic draft and its annotation revealed that *C. maltosa* has several hundred less genes than its pathogenic relatives *C. tropicalis* and *C. albicans*. Thorough computational scrutiny with alternative approaches strongly suggested that these are actual genes losses and not the result of problems in the genome assembly. The gene reduction does not seem to be specific to nonpathogenic species since *C. sojae*, a closely related nonvirulent species, also has considerably more genes than *C. maltosa*. In addition, the missing genes are not particularly enriched for genes related to pathogenicity. In fact, *C. maltosa* has orthologs of most of the genes that have been previously associated to virulence in *C. albicans*. Gene loss in yeasts has been associated to tolerance of varied stresses (Robinson et al., 2023; Steenwyk & Rokas, 2018), which could be the conditions that *C. maltosa* is encountering in the industrial environments where it has been isolated.

The case of the *TLO* genes is worth pointing out. These genes encode a conserved Med2 domain that is a component of the Mediator complex, with a possible role in the expression of genes related to virulence such as those involved in antifungal drug resistance and dimorphism (Flanagan et al., 2018; Haran et al., 2014). *C. albicans* has an unusually high number of TLO genes (15), while *C. maltosa*, *C. sojae*, and even the pathogenic *C. tropicalis*, have only one. *C. dubliniensis*, the closest species to *C. albicans*, also only has one ortholog suggesting that the large number of *TLOs* in *C. albicans* could represent gains associated to the ability of this species to colonize the human body, but that are not shared with other pathogenic species. Despite the specific cases, overall, the dissimilarities in virulence between these *Candida* species does not seem to be easily explained by broad changes in gene content. Instead, other evolutionary modifications such as those involving the regulation of genes may be more important underpinnings of these differences. Although further investigation is needed, the specific lower number of genes observed in *C. maltosa* seems related to its evolutionary history rather than to its inability to colonize the human body.

The auxotrophic strains that we generated here, together with previously available aminoacidic marker cassettes (Mancera et al., 2019), allowed the implementation of an effective strategy to genetically modify *C. maltosa*. Previously developed host-vector systems for this species relied on auxotrophic strain generated by random mutagenesis screens (Kawai et al., 1991; Nakazawa et al., 1997) with the risks of generating off-target mutations that these approaches have. In fact, high instability of some of these mutants has been observed (Kiss & Stephanopoulos, 1992). Taking advantage of the new genome assembly and a SAT-FLP cassette developed for *C. tropicalis*, we were able to specifically target three amino acid genes that have been typically employed to produce auxotrophies for the generation of gene deletion mutants in several related species. Two independent auxotrophic lines were generated so that deletions can be performed in replicates. Apart from using the triple auxotrophic strains for gene deletion as performed here, these strains could be the basis to implement CRISPR-Cas9 approaches in this species. For example, there is a CRISPR system developed for *C. albicans* that relies on the integration of the cassette in the *LEU2* gene of *C. maltosa* (Nguyen et al., 2017). This gene has been used as a complementation marker in large gene knockout efforts in *C. albicans* (Noble & Johnson, 2005). In principle, using the strains we have generated here, this system could be easily implemented to perform genetic modifications in *C. maltosa*.

As a proof of concept of the genetic modification approach, we successfully generated a homozygous gene knockout of the *EFG1* gene. In related CTG-species, this gene encodes a transcription regulator that controls several cellular transitions including filamentation, biofilm formation and white-opaque switching (Hirakawa et al., 2015; Mancera et al., 2015). All of these morphological processes have been associated to the subsistence of the pathogenic species in the human host and the balance between commensal and pathogenic states (Huangid et al., 2019; Montelongo-Jauregui et al., 2019). The involvement of *EFG1* in the regulation of some of these morphological transitions such as filamentation and biofilm formation, has been shown to be conserved as far as in *C. parapsilosis*. In agreement, disruption of *EFG1* in *C. maltosa* led to phenotypic changes in filamentation and biofilm formation. Interestingly, contrary to the observation in *C. tropicalis*, the *efg1* mutant showed increased filamentation capacity in the conditions here employed. These results agree with previous observations in *C. parapsilosi*, in which Efg1 could be operating as a negative regulator of filamentation (Connolly et al., 2013). Our results also suggested that the defect in biofilm formation of the *C. maltosa efg1* mutant does not fully depend on a negative effect in filamentation.

Despite the conservation in the involvement of *EFG1* in these morphological transitions, what is clearly different is the reduced ability of *C. maltosa* to filament and form biofilms as the ones formed by the pathogenic species (Mancera et al., 2021). In the *in vitro* assay that we have routinely used, the biofilms formed by *C. maltosa* are considerably lighter and less structured. It is possible that this species can filament or form biofilms under different conditions, although we did test several media some of which resemble the conditions where *C. maltosa* has been isolated. These observations suggest that *C. maltosa* is overall a poorer filament and biofilm former or that it establishes these morphological structures under conditions that are considerably different than those where the pathogenic species form biofilms.

Research involving *C. maltosa* has deaccelerated in the last ten to fifteen years; the focus that this species received by research groups mainly in Japan and Europe during the nineties and at the beginning of the current century (Mauersberger et al., 1996; Takaku et al., 2001, 2002) seems to have dissipated. We propose using this species as a comparative model to better understand the molecular basis of the virulence of closely related *Candida* species, especially *C. tropicalis*. The genomic draft and genetic modification strategies presented here will allow such molecular comparisons and have the potential to facilitate research towards biotechnological applications with *C. maltosa*.

## DATA AVAILABILITY

The raw genome data, the genome assembly and the annotations have been deposited the NCBI under BioProject ID PRJNA1036158.

## ACKNOWLEDGEMENTS

We thank Susana Ruiz-Castro, Maria Sofia Pardo Reyes, and Claudia Geraldin León-Ramírez for technical assistance. This work was funded by Consejo Nacional de Ciencia y Tecnología de México (FORDECYT-PRONACES/103000/2020); MACT and LFG-O were funded by Conacyt at the doctoral (2019-000037-02NACF-26396) and postdoctoral (4133922) level, respectively. The funders had no role in study design, data collection and analysis, decision to publish, or preparation of the manuscript.

## MANUSCRIPT VERSIONS

This manuscript was released as a pre-print at *bioRxiv* (Chávez-Tinoco M., et al., 2023).

## TABLE LEGENDS

Supplementary Table 1. Lists of pathogenicity and filamentation associated genes and their orthologs.

Supplementary Table 2. Primers used for the genetic modification of *C. maltosa*.

